# Stacks 2: Analytical Methods for Paired-end Sequencing Improve RADseq-based Population Genomics

**DOI:** 10.1101/615385

**Authors:** Nicolas C. Rochette, Angel G. Rivera-Colón, Julian M. Catchen

## Abstract

For half a century population genetics studies have put type II restriction endonucleases to work. Now, coupled with massively-parallel, short-read sequencing, the family of RAD protocols that wields these enzymes has generated vast genetic knowledge from the natural world. Here we describe the first software capable of using paired-end sequencing to derive short contigs from *de novo* RAD data natively. Stacks version 2 employs a *de Bruijn* graph assembler to build contigs from paired-end reads and overlap those contigs with the corresponding single-end loci. The new architecture allows all the individuals in a meta population to be considered at the same time as each RAD locus is processed. This enables a Bayesian genotype caller to provide precise SNPs, and a robust algorithm to phase those SNPs into long haplotypes – generating RAD loci that are 400-800bp in length. To prove its recall and precision, we test the software with simulated data and compare reference-aligned and *de novo* analyses of three empirical datasets. We show that the latest version of Stacks is highly accurate and outperforms other software in assembling and genotyping paired-end *de novo* datasets.

## Introduction

Type II restriction enzymes [Smith1970; Kelly1970] remain one of the primary drivers in population genetics experiments. Starting with their first application in the mid-1970s [Grodzicker1974; Botstein1980], restriction enzymes have been paired with advancing technologies, including the polymerase chain reaction coupled with polyacrylamide gel analysis [Blears1998], microarrays [Miller2007], and most recently, massively parallel, short-read sequencing to yield great insights into model and natural populations. While sequencing costs have decreased by orders of magnitude since the completion of the human genome project, the cost is still too high in the majority of ecologically-based, non-model studies for whole genome re-sequencing, leaving a wide niche for the set of Restriction-site Associated DNA sequencing (RADseq) protocols.

RADseq has grown into a family of protocols whose kin have been optimized to different criteria. The protocols vary on a few major axes; first, a protocol may employ one (GBS, single-digest RAD) or two restriction enzymes (CRoPS, double-digest RAD, ddRAD). Second, it may rely on the distance between cut sites to determine the length of DNA that is sampled (CRoPS, GBS, ddRAD), or it may employ sonication to create relatively uniform lengths of DNA (sdRAD, BestRAD). Third, protocols may use a size selection step to explicitly select the length range of DNA molecules to enrich, or they may rely on PCR to enrich shorter sequences for the final sequencing library. Additional protocols are further specialized, focused on adapting more of the steps to off-the-shelf kits (EasyRAD). Others were designed to minimize primer-dimers (3RAD [Graham2015]), or to use type IIb restriction enzymes (2bRAD [Wang2012]), or to use biotinylated adapters to extract restriction site-associated DNA from other genomic DNA (BestRAD [Ali2016]), as well as hybrids of the above approaches (see [Davey2011] and [Andrews2016] for reviews).

While RADseq can produce orders of magnitude more genetic markers than earlier marker technologies, WGS produces an order of magnitude greater still. The primary obstacle, however, remains cost. It is popular – but misguided – to relate changes in sequencing technologies to Moore’s Law; Moore’s Law requires a halving of cost every 18 months and has held for sixty years, while sequencing technology has instead reduced cost by orders of magnitude twice (first with 454 pyrosequencing, and later with Illumina sequencing-by-synthesis) [http://genome.gov/sequencingcosts], and while Illumina has continued to reduce costs incrementally, there is no clear path to any order-of-magnitude-changes on the horizon. In fact, long molecule sequencing is significantly more expensive than technology that came prior. For every 560Mb threespine stickleback fish genome that is WGS sequenced, 120 genomes can be examined with RAD for the same sequencing cost and with a single library preparation (WGS requires a library per genome). Nothing in the near future is yet predicted to change this resource advantage for large studies.

The union of genome sampling protocols with massively parallel, short-read sequencing has produced an immensely successful research program in population [Bassham2018], conservation [Dierickx2015] and landscape genomics [Bay2018], phylogenetics [Spriggs2019], and epigenetics [Trucchi2016], creating new experimental space for non-model organisms, and allowing, for example, ambitious sampling regimes in large geographical surveys [Dudaniec2017], as well as wide-ranging taxon breadth in phylogenetic studies [Near2018]. Regardless of the analytical approach, and in addition to any challenges of the experimental design, all RADseq strategies present two fundamental issues. First, the precision of the analytical results depends significantly on the quality and amount of DNA available. RADseq has expanded the scope of organisms that can be examined, but sampling many of these organisms from nature is a challenge and, in these cases, DNA may be degraded or available only in small quantities. Second, restriction sites may not be conserved across all individuals in the experiment, depending on evolutionary distance between them, and the length of the restriction site(s) of interest. In both cases, the molecular library may not contain enough template molecules to have sufficiently sampled all of the alleles in the genome in each individual. Two additional processes will sample these template molecules in the library – PCR amplification will sample molecules to create additional copies, and the sequencer will select from the amplified molecules for inclusion on the sequencing flow cell. Having too few templates for amplification or selecting too few molecules to sequence (a low depth of coverage) can exacerbate the effects of allelic dropout.

Through our participation in a large number of studies conducted over the last decade, both our own, collaborations in the field, and by interacting with scores of scientists in the support of Stacks version 1 (v1), we have learned a lot. The quality of DNA, the success of library preparation, and the sequencing strategy – all contributing to differential allelic sampling – can separate the path-breaking RADseq studies from the rest. Often, the differences between these studies generated substantial discussion in the community [Lowry2017; McKinney2017; Catchen2017] and a lot of speculation as to the inherent limitations of reduced representation sequencing.

Our experience is that with the development of proper analytical protocols [Paris2017; Rochette2017], and with supportive software, we can close the performance gap between the path-breaking RAD studies and the rest and secure a successful experimental strategy for the next decade. We sought to design a software that could help identify poorly performing libraries and provide support in the design of new studies. We sought to maximize the amount of information we could extract from RAD protocols by focusing on Illumina paired-end, short-read sequencing and improving the analysis tools to provide the most polymorphic loci possible and the richest set of haplotypes to increase information yield significantly.

The collection of software to implement this strategy has resulted in the second major version of Stacks. Version 2 (v2) incorporates paired-end reads natively into the locus assembly algorithm providing for locus sizes greater than 500bp, increasing the number of polymorphic loci, significantly improving genotyping accuracy, and providing phased haplotype markers for those loci, in a massively scalable form. As we show, v2 outperforms every other RAD software. Finally, to vet and optimize the software, we designed and implemented an accurate RAD simulation system which shed light on basic processes, like the effects of PCR duplicates and sequencing coverage while providing us with a road map to fully optimize our algorithms.

## Methods

### Changes to the Stacks pipeline

Stacks v1 was designed to process individual samples independently, identifying polymorphic sites within each individual (`ustacks` or `pstacks`), then connecting loci across samples, through the creation of a catalog, to provide a single view of the meta-population (`cstacks`). Individuals could then be matched to the metapopulation data contained in the catalog with `sstacks` [Catchen2011]. Availability of computational resources undergirded this design decision as the pipeline needed to process potentially thousands of individual samples, each with millions of raw reads. This architecture prevented population-level information, such as presence of a polymorphic site, from being incorporated into individual genotype calls. To incorporate this population-level information, version 1.10 of Stacks (2013) incorporated the `rxstacks` program to make population-level corrections retrospectively, after the core pipeline had executed. The major architectural change to the Stacks v1 pipeline is the reorganization of individual-level data so that it is stored per locus instead of per-individual. This cosmetically simple change, implemented in the tsv2bam program, enables large analytical gains downstream in the pipeline (Fig. 1).

**Figure 1.**
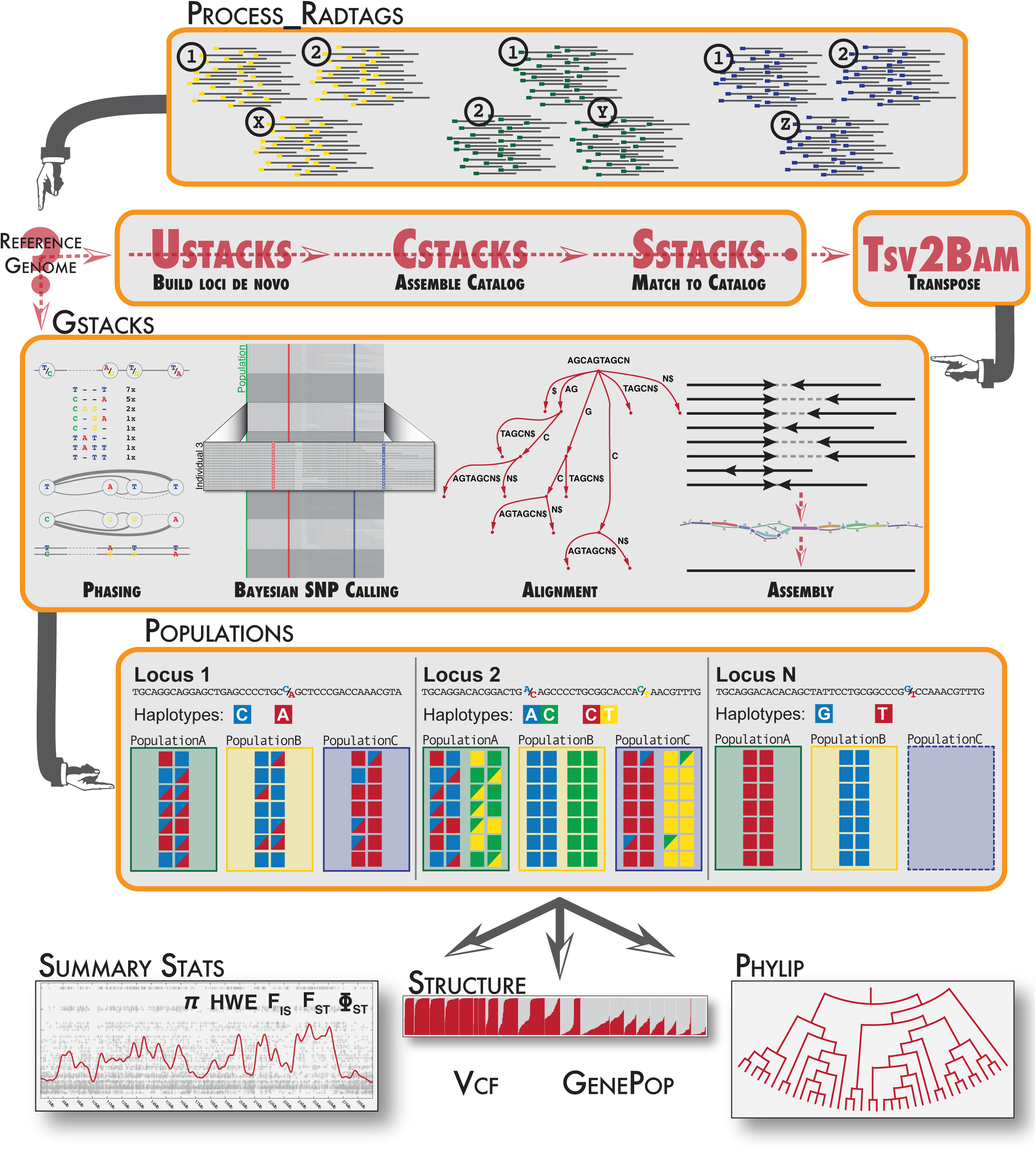
Stacks v2 pipeline overview.

The v2 pipeline for *de novo* analyses starts by clustering loci as in Stacks v1 using `ustacks`, `cstacks` and `sstacks`, but then continues with tsv2bam and additionally with the `gstacks` program, which now forms the analytical core of the v2 pipeline. The `populations` program, which applies a population genetic-frame to the data, allowing for advanced data filtering and data export, has also been rearchitected to process loci in a streaming fashion – locus by locus – where previously all loci from all samples had to fit into computer memory at once, leading to a memory-bound program.

In sum, the new *de novo* architecture is simpler, more versatile and substantially faster than Stacks v1’s `rxstacks`-`cstacks`-`sstacks`-`populations` sequence. For reference-based analyses, the main Stacks pipeline now begins directly with the `gstacks` program (and the v1 clustering pipeline is not employed) followed by a call to the `populations` program.

### Improved locus clustering procedure

For *de novo* analyses, the core of the Stacks clustering algorithm (`ustacks`-`cstacks`-`sstacks`), which builds loci out of the single-end reads, remains as previously described [Catchen2011, Catchen2013], but has received a number of improvements and optimizations. Stacks has been capable of gapped assemblies since version 1.38 (2016), when Needleman-Wunch comparisons between stacks sharing many k-mers were added, and in Stacks v2 this capability has been refined and has become the default. In addition, while specific parameter optimization is still necessary [Paris2017, Rochette2017], several assembly parameters have had their default values changed in an attempt to offer better initial results for a wider range of datasets.

### Transposing sequence data storage

The new tsv2bam program, which concludes the locus clustering stage, sorts the reads (or read pairs) of each individual by catalog locus. Paired-end reads can be incorporated by matching the set of single-end read IDs in each locus. This is similar in principle to the sorting of alignment files by chromosome and coordinate, a strategy employed by most reference-based analysis pipelines. Importantly, this sorting step allows Stacks to stream the data locus by locus in subsequent analyses in `gstacks`, making it computationally feasible to work, at each locus, with the full sequencing information for all samples simultaneously. This enables `gstacks` to assemble a contig, align all reads from the meta-population to that locus, call polymorphisms across the population, and call population-informed genotypes in each individual sample.

### RAD-locus contig assembly

In *de novo* mode, `gstacks` starts by assembling a short contig for each locus (i.e. for the set of single-end reads clustered by `ustacks`-`cstacks`-`sstacks` across the metapopulation and the associated paired-end reads fetched by tsv2bam). The reads of the locus (by default up to 1,000 reads or read pairs sampled uniformly across individuals) are broken into k-mers (by default with k=31) and inserted into a *de Bruijn* graph (Fig. 2). K-mers with a coverage lower than a minimum threshold (by default 2) are discarded. Stacks v2 then scores each connected subgraph for total coverage from forward and reverse reads (if any). The subgraphs with the highest total coverage for forward and reverse reads are extracted (they may be confounded); other subgraphs are discarded. If the subgraph for reverse reads is not a directed acyclic graph (a graph with no loops), reverse reads are discarded and the graph is recomputed using forward reads only. If the subgraph for forward reads is not a directed acyclic graph, the entire locus is discarded. Some elements of DNA, such as microsatellites, make very small and predictable loops in a *de Bruijn* graph. We screen for such elements and clip the k-mer(s) linking the microsatellite repeat to remove these simple loops and prevent the subgraph from being discarded.

**Figure 2.**
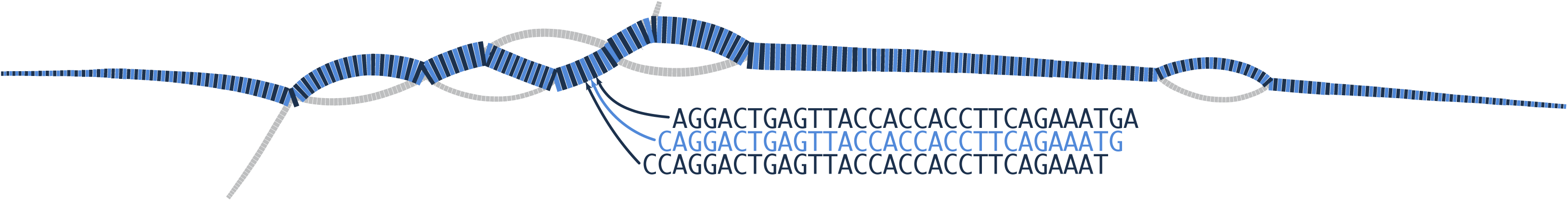
Locus assembly and read alignments.

The path in the ‘forward’ subgraph that has the highest total coverage is identified using an exact depth-first algorithm. Using highest coverage as the scoring criterion favors the inclusion of major alleles over minor ones and marginally of insertions over deletions. This path is converted back into a nucleotide sequence.

If there are no reverse reads or if the ‘reverse’ subgraph is confounded with the ‘forward’ one, the ‘forward’ sequence is the reference contig for the locus. Otherwise, if there is a distinct ‘reverse’ subgraph, the above procedure is repeated to obtain a ‘reverse’ sequence and overlapping between the ‘forward’ and ‘reverse’ contig sequences is attempted.

To overlap the single-end (SE) contig with the assembled, paired-end (PE) contig a suffix tree is constructed from the PE contig using Ukkonen’s algorithm [Gusfield1997, chapter 6], which can build the tree in linear time. The consensus sequence from the SE locus is then aligned against the suffix tree in linear time and the set of maximal matches is recorded. If a perfect match against the suffix tree is recorded, then the SE and PE sequences are merged and the operation is complete. However, if there are two or more fragment matches between the SE and PE contig, the set of alignment fragments are converted into a directed, acyclic graph (DAG) to find the best set of non-overlapping alignment fragments. A Needleman-Wunsch gapped alignment matrix is then initialized and populated with the set of alignment fragments. Finally, a bounded, gapped alignment is performed – looking only at the regions of the alignment matrix that were outlined by the suffix tree alignment fragments – to complete the overlap between the SE locus and PE contig. If no matches were found against the suffix tree, the 3’ end of the SE locus is compared directly against the 5’ end of the PE contig using a Needleman-Wunsch gapped alignment to check for any overlap that is below the minimum match length of the suffix tree. If there is no overlap between the forward and reverse contig segments, the two sequences are merged into a single scaffold with ten ‘N’ characters, to symbolize a gap of unknown length between the forward and reverse regions.

The algorithms described here also function well for paired-end data generated from two-enzyme double-digest protocols such as ddRAD. The forward reads (*i.e.* those starting with the restriction site that was ligated to the forward sequencing adapter) are first used to cluster read pairs into RAD loci, then the forward and paired-end reads are used to assemble a contig for each locus. However, since both the forward and reverse reads are anchored to a restriction site, coverage will be uniform on both sides of the locus, and the length of the contig will reflect the properties of the underlying genomic sequence. If the two restriction sites are less than two read lengths apart, Stacks will overlap the two sides and yield a single continuous contig, otherwise it will produce a scaffold comprising two contigs (one for each side). There exist limit cases where the distance between restriction sites varies within the population due to structural variation or restriction site polymorphism and rescue. For example, if the second enzyme cuts frequently there may be multiple restriction sites nearby. If those sites are variable within the population, different individuals may return paired-end reads anchored to different restriction enzyme cut sites. The algorithm handles these cases naturally and will report contigs faithful to the underlying sequences but that will often have atypical properties, such as being both continuous and longer than two read lengths.

Lastly, we note that the present approach is not applicable to the GBS protocol, which uses a single-enzyme double-digest strategy, because the restriction fragments and subsequently the sequencing library inserts intrinsically lack an orientation, so that it is not possible to single out half the reads for locus clustering.

### Read alignments

Once the full locus has been assembled, yielding a consensus sequence, individual reads must be aligned in a per-sample context. This operation proceeds in a methodologically similar way to the contig overlap stage: a suffix tree is built from the locus consensus sequence, which was combined from the SE locus and the PE contig. Reads are then aligned against the suffix tree, with three possible outcomes. First, no alignments may be found, in which case the read is ignored. Second, a perfect alignment – encapsulating the full length of the read with no mismatches — may have been found against the suffix tree, from which an alignment CIGAR is calculated and recorded, or third, more than one maximal match was found against the suffix tree in which case the alignment fragments are ordered into a DAG, the consistent alignments from the DAG are used to populate a Needleman-Wunsch gapped alignment matrix, and a bounded, gapped alignment is conducted to connect the aligned fragments for the final alignment. A CIGAR string representing the alignment is recorded and the process is repeated until all reads from all samples have been aligned.

### Reference-based analyses

In *reference mode*, Stacks v2 begins directly with the `gstacks` program (Fig. 1). Alignment of RADseq reads is done by an external alignment program (e.g. BWA [Li2009b, Li2013] or Bowtie [Langmead2012]) and `gstacks` expects to have a set of aligned and reference-sorted reads for a series of samples, typically provided as one sample per BAM file [Li2009a]. The `gstacks` program uses a sliding window strategy to build RAD loci from sets of alignments. Starting at the first sample, `gstacks` identifies the 5’ alignment position of the first forward read and uses this basepair position to anchor the sliding window. Assuming the first read (and the RAD cutsite) is on the positive strand, reads that are aligned within the length of the window are added to the locus if they share the same 5’ alignment position. If reads within the window are on the negative strand, their 5’ starting positions are calculated from the CIGAR string and they are also added to the locus. Reading completes for the first sample and locus when the boundary of the window is reached (by default, 1Kbp in length). The `gstacks` program them iterates over the remaining samples for the same window position, collecting all read alignments. After the data has been acquired from all samples for the current locus, the locus is stored and the window is advanced to the next read beyond the previous window bound. If an initial read filling the window is a paired-end read, then the RAD cut-site is on the negative strand (and the starting basepair is on the 3’ end of the RAD locus), and the locus starting position is calculated by adding the length derived from the alignment CIGAR string to the 5’ alignment position. Once a locus has been built from the aligned reads across the meta-population, the same filtering and genotyping methods are applied as in the *de novo* mode.

### PCR duplicates removal

For paired-end data derived from shearing-based protocols, PCR duplicates can be removed by identifying inserts that map to the same start and end coordinates and randomly discarding all but one of them. This approach relies on the randomness of the shearing process, which creates a mixture of inserts of diverse lengths, that are then preserved through PCR amplification. Distinct source molecules may have identical insert sizes by chance, but this is rare for RADseq data, assuming the sequencing depth (e.g. 30X) is less than the width of the insert size window (e.g. a 100bp window, with inserts ranging from 300 to 400bp), so discarding inserts of identical sizes efficiently and selectively removes PCR duplicates. In the `gstacks` program, in both *de novo* and reference mode, after the locus has been assembled and reads aligned (or the locus has been populated from the pre-aligned reads), for each locus, reads are sorted by sample and then by insert length, and all reads within one individual with the same insert length are discarded save one pair.

If reads are derived from a ddRAD protocol, then PCR duplicates cannot be detected by `gstacks`, because differential insert lengths cannot be used as a means to detect them (since all reads will be attached to one of the two restriction enzyme cutsites). If a protocol which incorporates a random oligo sequence into the barcode is used for construction, then PCR duplicates could still be detected, prior to running the main pipeline, using the `clone_filter` program of Stacks.

### Genotyping model

In Stacks v1 we employed a SNP calling method first proposed in [Hohenlohe2010] which itself was modified from [Lynch2009] to allow for a per-nucleotide sequencing error rate to be estimated. This model worked well, however, it required relatively high coverage for robust results and it could not incorporate population-level data, since it only had access to sequence reads from one individual at a time. For Stacks v2, we again incorporated the work of Lynch and colleagues, who have continued to improve their SNP calling method by adding a Bayesian genotype caller (BGC) [Maruki2017]. In this framework, the `gstacks` program will identify the presence of a SNP within a locus by examining the read data from the entire metapopulation. This support for the presence of a SNP is then fed into the genotyping model as a Bayesian prior, which incorporates the metapopulation information to genotype each individual separately at that SNP position. The BGC model does not require any multiple testing corrections, however, it is designed for diploid genomes and expects two alleles per site. Stacks v2 implements several alternative models to call SNPs and genotypes: BGC (Bayesian genotype caller), the default model just described, along with the HGC (high-coverage genotype caller) [Maruki2017], and we still provide the method of [Hohenlohe2010] (which was the default in Stacks v1). For the BGC model, the following numerical stabilizations were used: (i) when computing the sequencing error rate under the assumption of polymorphism (equation 6 in [Maruki2015]) we always assume at least 0.1 error nucleotides have been observed across the population; (ii) the genotype frequencies used in genotype likelihood computations are rescaled so as to always be greater or equal to 1 over the number of samples.

### Converting SNPs into phased haplotypes

SNP alleles that are observed on the same read or within the same read pair are part of the same haplotype, because a read or read pair (or library insert) is a sequence sample from a specific chromosome. We can take advantage of this natural phasing to provide sets of haplotypes from each RAD locus. While Stacks v1 provided short haplotypes encompassing the single-end locus, Stacks v2 extends this functionality by phasing heterozygotes using a graph-based algorithm that relies on sequence data, specifically on co-observations of alleles within a read (or read pair).

After genotyping, the `gstacks` program attempts to build locus-wide haplotypes for each individual. In other words, if a locus includes several SNPs and an individual is heterozygous at two or more of them, Stacks v2 reconstructs the combination of alleles that corresponds to the individual’s two chromosomes (Fig. 3). To this end, Stacks v2 implements a read-based phasing approach (as opposed to statistical phasing [Browning2011]). Essentially, this approach relies on the co-observation, in a given read (or read pair), of the alleles at several SNPs. A graph is built in which the nodes are the alternate alleles at heterozygous SNPs, and the edges are the co-observations of these alleles in reads. Given that one read can only have a single nucleotide at a given position, edges always link alleles of distinct SNPs. To limit the influence of sequencing errors, edges with a weight (i.e. number of co-observations, or coverage) less than a given threshold (by default 2, alleles seen together only once) are ignored. If there are no conflicts between reads, the haplotype graph will be composed of two or more distinct connected subgraphs and the corresponding haplotypes can be extracted from each subgraph. Otherwise (*i.e.* if there is substantial conflict between reads) the phasing operation was likely confounded by sequencing errors, contamination, or over-merging. In this case, the individual’s SNP alleles are marked as unreliable, as they are likely affected by the same issues, and no phasing is provided.

**Figure 3.**
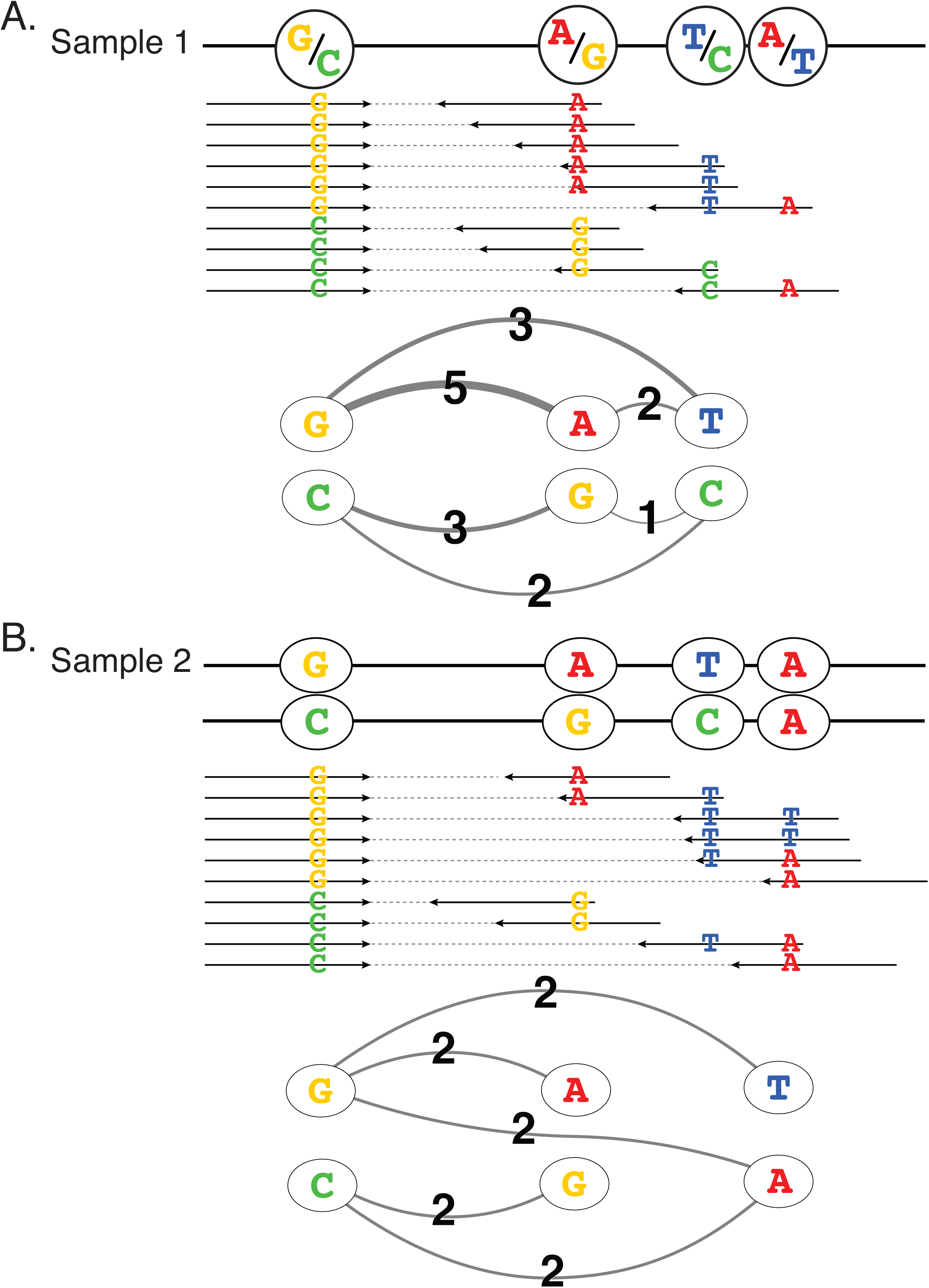
Algorithm for haplotyping heterozygotes. (A) Sample 1 is successfully phased at this locus as all alleles are found to co-occur on two or more read pairs producing two distinct subgraphs. (B) Sample 2 cannot be phased as at the fourth variable position the ‘A’ allele is observed on two different allelic background, confounding the haplotype graph.

### Gstacks output files

The `gstacks` program proceeds through the data set one locus at a time and can be parallelized to run one locus per thread at a time. The program will produce a new catalog contained in two files: the consensus sequences for the catalog loci in a FASTA formatted file, and a custom, compressed file containing all of the SNP/haplotype calls for each locus and all individuals. These files are designed to be read by the `populations` program, which can filter the loci and export them. In addition, `gstacks` provides a number of useful data distributions, such as coverage and phasing rates, in its output.

### Improvements to the populations program

The `populations` program is designed to take the assembled data from `gstacks` and apply a population genetics frame to the data. The program can apply a population map to the underlying data to segment the individuals in to populations based on useful criteria, like geography, phenotype, sex, or another category. The program can filter data in a number of useful ways: keeping polymorphic sites that are found within a specified number of individuals or populations. A new option allows samples to be filtered according to haplotype presence. A user can ask to keep all haplotypes that are found (and complete) in, for example, more than 80% of individuals. Remarkably, for a particular locus, the set of haplotypes will be constructed from a number of SNPs, but even if all SNPs are present in 80% of individuals, it is possible that different SNPs will be missing in different individuals, resulting in less than 80% of complete haplotypes. In order to provide the longest possible haplotype at the required threshold, the `populations` program will filter individual SNPs by their availability in the population until the criterion is met.

After filtering, the `populations` program will calculate a number of population genetic statistics, for both SNPs and haplotypes. These include π, heterozygosity, F_IS_ computed per-population and per-SNP, as well as F_ST_ computed for each pair of populations using an Analysis of Molecular Variance (AMOVA) approach [Weir1996; Weir1984; Excoffier1992]. For haplotypes, several versions of F_ST_ are calculated, including φ_ST_ [Holsinger2009; Bird2011] and F_ST_’ [Meirmans2006]. The `populations` program also provides a measure of Hardy-Weinberg equilibrium for each SNP [Engels2009; Louis1987] and each locus [Guo1992]. The SNP-level (and therefore two-allele) approach uses an exact test that is analogous to Fisher’s exact test. For haplotypes, where multiple alleles can make an exact test computationally challenging, we employ a Markov chain approximation as described by Guo and Thompson (1992).

The `populations` program is able to export data, after filtering, in a number of useful formats from VCF to FASTA, and for specialized programs such as STRUCTURE, PLINK, or GenePop.

### Real datasets

We examined three empirical datasets to assess the capabilities of Stacks v2 and to compare it with software systems in the same class. First, we looked at 241 North American yellow warbler individuals (*Setophaga petechia*) contained in 18 populations from [Bay2018]. The 2×100bp RADseq data were generated with the BestRAD protocol [Ali2016] using the *SbfI* 8bp restriction enzyme. For reference-based analyses, we used the scaffold-level yellow warbler reference genome [Bay2018] which is assembled to 18,144 scaffolds with a length of 1.26Gbp and an N50 of 491,655bp. Second, we looked at 10 threespine stickleback (*Gasterosteus aculeatus*) individuals from two populations near Cook Inlet, Alaska [Nelson2018]. The RAD library derived from these fish was generated using the single-digest, sheared RAD protocol [Baird2008] with a 6bp *PstI* restriction enzyme and sequenced to 2×250bp. For reference analysis, the data were mapped to the threespine stickleback reference genome (BROADS1, Ensembl version 84). Finally, we examined 71 Antarctic bald notothen individuals (*Pagothenia borchgrevinki*) in two populations from McMurdo Sound and Prydz Bay, Antarctica (unpublished). The RAD library generated used the single-digest, sheared RAD [Baird2008] with the 8bp *SbfI* restriction enzyme. For reference-based analyses, we used a draft assembly of *P. borchgrevinki* (unpublished), which has a length of 0.76Gbp derived from 9399 scaffolds, with an 726,105bp N50.

### Simulations

We simulated paired-end RAD reads using the *RADinitio* software pipeline, version 1.0 [Rivera-Colón et al., in prep.]. The *RADinitio* software uses *msprime* [Kelleher2016] to generate coalescent trees for various samples given a user-defined neutral demographic model that can then be converted into individual haplotypes. We used the threespine stickleback (*Gasterosteus aculeatus*) reference genome (Ensembl version 84) as a base for our simulations, evolving four populations each with an effective population size of 10,000 haploid individuals, of which we sampled 25 diploid individuals per-population. The simulations had a mutation rate of 5 × 10^−8^ per base and per generation, a symmetric migration rate of 0.05 chromosomes per-generation, a flat recombination rate of 100cM per chromosome and an indel-over-substitution ratio of 0.1 (9% of mutation events will be short indels). *RADinitio* finds natural RAD loci via an *in silico* digestion of the evolved population of genomes. We simulated a single-digest library preparation [Baird2008; Etter2011] using the *SbfI* restriction enzyme and extracted the RAD loci. Sequencing is then simulated over the extracted RAD loci with a specified target sequencing coverage (10 or 20x), a read length of 144bp, an insert size sampled from a normal distribution (mean 350bp, standard deviation 70bp), and a sequencing error rate increasing linearly along the 5’ to 3’ length of the read from 0.1% to 0.5% over 144 base pairs.

For reference-based approaches, simulated reads were mapped to the stickleback reference genome using the *mem* routine of *BWA* [Li2013]. For *de novo* approaches, the consensus sequences of the reconstructed loci were mapped back to the reference genome using the same process.

We computed recall (fraction of the simulated SNPs/genotypes that were discovered) and precision (fraction of calls that were correct) by comparing the genotypes obtain via Stacks v2 against the reference genotypes obtained directly from the *RADinitio* simulations using custom Python scripts. We smoothed the results by analyzing 10 replicates of each simulated dataset.

### Result filtering for comparison and evaluation

To evaluate Stacks v2 and to make reasonable comparisons to other software we have to select a rational subset of the simulated and empirical RAD data for comparison. There is significant variation in the reconstruction of loci that is unrelated to the performance of the analytical software. For example, since we are testing sheared, single-digest protocols, the length of the locus ‘sequenced’ *in silico* will vary, and so the 3’ region of the locus will not be precisely defined, and this region will include polymorphic sites. An exact comparison to the simulated data is not therefore possible. Likewise, in a *de novo* assembly of empirical data, the number RAD loci reconstructed will be highly biased towards a very large number of loci assembled in just one individual (see [Rochette2017], Fig. 3 for details), thus heavily biasing any assessment of results.

For that reason, when discussing the number of loci reconstructed, we report loci found in 80% of individual samples in each data set (NS80 loci). We do so for both simulated and empirical data sets. When discussing SNPs, we report SNPs that have a minor allele count of at least 3 (MAC3). Given our diploid data sets, this ensures that a SNP has been found in at least two individuals. We report genotypes as those that could be called in 80% of individuals (GT80). Finally, when discussing assignment of haplotypes at a locus, we report the number of SNPs phased into haplotype that are found in 80% of individuals (the raw number of haplotypes is equal to the number of polymorphic loci). To find haplotypes (see ‘populations’ section above) that are fully phased in all individuals (no unknown sites), some SNPs will be filtered, and a lower value is reported here than when looking at the number or independent GT80 sites. Finally, for simulations, we report the number of *true* SNPs that were found that are both MAC3 and NS80 (TNS80). All of the above values are listed in Table 1.

**Table 1.**
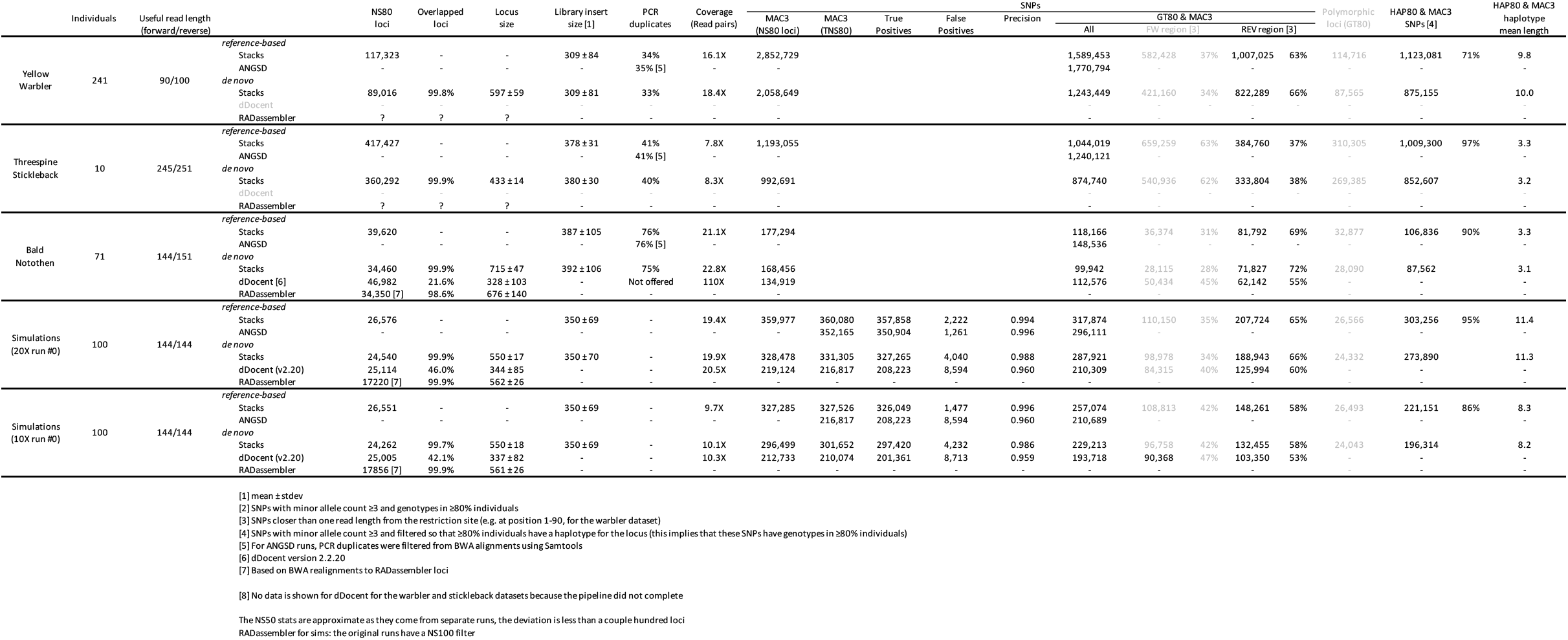
General statistics for all datasets & approaches.

## Results

### Calling SNPs, genotypes and haplotypes from paired-end RADseq data

We first surveyed the general properties of paired-end data derived from single-digest, shearing-based RADseq protocols. Using a reference genome approach, we reanalyzed three published datasets, including the Alaskan threespine stickleback [Nelson2018] (sdRAD, PstI enzyme, 2×250bp reads), the North American yellow warbler [Bay2018] (BestRAD, SbfI enzyme, 2×100bp reads), plus one newly generated dataset for a teleost fish, the Antarctic bald notothen (*Pagothenia borchgrevinki*) (sdRAD, SbfI enzyme, 2×150bp reads).

We aligned the paired reads of all individuals to their respective reference genomes using BWA [Li2009b, Li2013], then used Stacks to identify RAD loci, remove PCR duplicates and compute the distribution of insert lengths (Fig. 4, A-C, solid lines) as well as the per-basepair coverage across the locus (Fig. 4, A-C, grey shaded area). We found the median insert sizes for warbler to be 300bp, for stickleback to be 380bp, and for *P. borchgrevinki* to be 300bp, respectively. These distributions reflect the DNA shearing and size-selection steps of the library preparation. We examined the *in vitro* insert length of *P. borchgrevinki* using the Applied Biosystems 3730XL fragment analyzer and found that, after accounting for the additional length of the sequencing adaptors, Stacks v2 was accurately reconstructing the observed set of insert lengths (Fig. S1).

**Figure 4.**
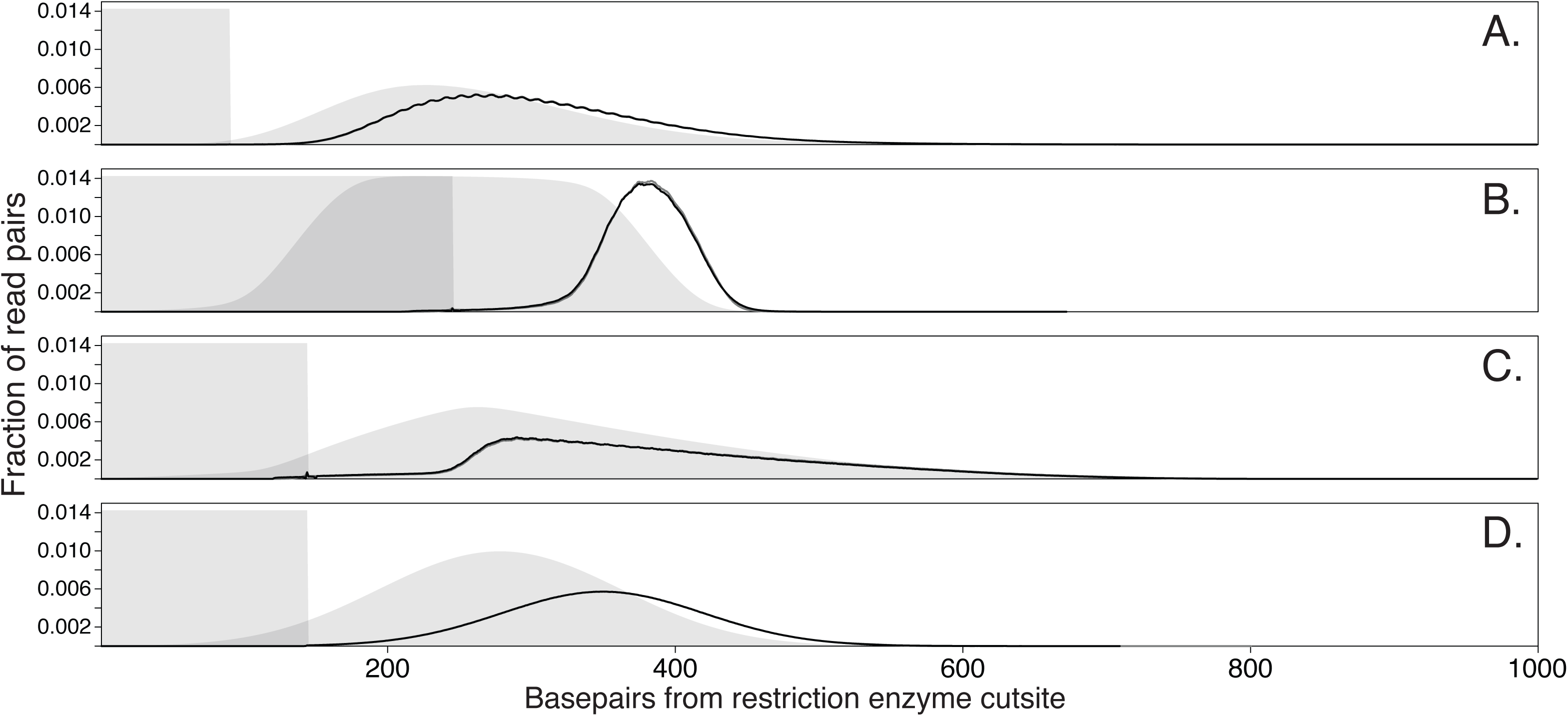
DNA library insert size distributions in the benchmark paired-end sdRAD datasets (A-E) as reconstructed by the reference-based (black lines) and *de novo* (gray lines) approaches. The shared areas represent the expected variation of the sequencing coverage along the length of RAD loci, based on read length of each dataset (respectively 100, 250, 150 and 150; see Table 1) and the insert size distribution. Expected coverage is directly derived from the insert size distribution, after all inserts are stacked by placing the first base of the restriction site at position 1. (A) Yellow warbler, (B) threespine stickleback, (C) bald notothen, (D) simulations, 20x. The periodic pattern of insert sizes apparent for the warbler dataset, and to some extent for the bald notothen one, seems to be real (Fig. S8).

In the case of sheared-RADseq data, in which inserts are anchored on one side by the restriction site, the distribution of insert sizes has direct implications for the distribution of sequencing coverage within each locus. Indeed, the principle of paired-end sequencing is to sequence both ends of each insert; for sheared-RADseq this leads to the coverage patterns shown in Figure 4 (A-C), where coverage on the restriction site end is constant over one read length, whereas coverage on the sheared end has a trapezium-like distribution (which appears triangle-like, or bell-like, if the width of the insert size distribution is larger than the read length). Figure 4 makes clear that three different size selection strategies were taken with the construction of the different libraries. While the stickleback library focused on a narrow size range while employing long Illumina reads – presumably to achieve uniform coverage – the *P. borchgrevinki* library focused on a wider insert length – presumably to assemble larger contigs – with the warbler dataset in between.

In turn, coverage affects polymorphism discovery and genotype calling over the length of each locus. SNP discovery remains efficient even at low coverage (Fig. 5A-C) because in Stacks v2 the existence of a polymorphism is a test on the population: evidence for alleles can be aggregated across individuals, so alternate alleles are visible as long as their total coverage in the population is substantial, independently of each individual’s coverage [Maruki2015]. Genotype calls, in contrast, fundamentally relate to single individuals, and are therefore much more affected by coverage variations (Fig. 5E-G; Fig. S2). Ultimately, whether genotypes can be called depends on the total coverage of the locus and on the dispersion of paired-end read positions. For the stickleback, 1.04 million SNPs were found that could be genotyped in ≥80% of individuals, with 37% found in the paired-end region (Table 1). In warbler, 1.59 million SNPs in the same class were found (63% in the paired-end region), and in *P. borchgrevinki*, 0.12 million SNPs were likewise found, with 69% in the paired-end region. The distribution of SNPs across the single- and paired-end regions in the three datasets reflects the insert length of the underlying RAD libraries and the natural polymorphism rate. In the case of the bald notothen, there are relatively few RAD loci, but two-thirds of the SNPs were found with the addition of the paired-end contig greatly increasing the value of the loci sampled by the RAD protocol (by returning more polymorphic loci). These results demonstrate that coverage from paired-end reads can effectively be turned into SNPs and genotypes.

**Figure 5.**
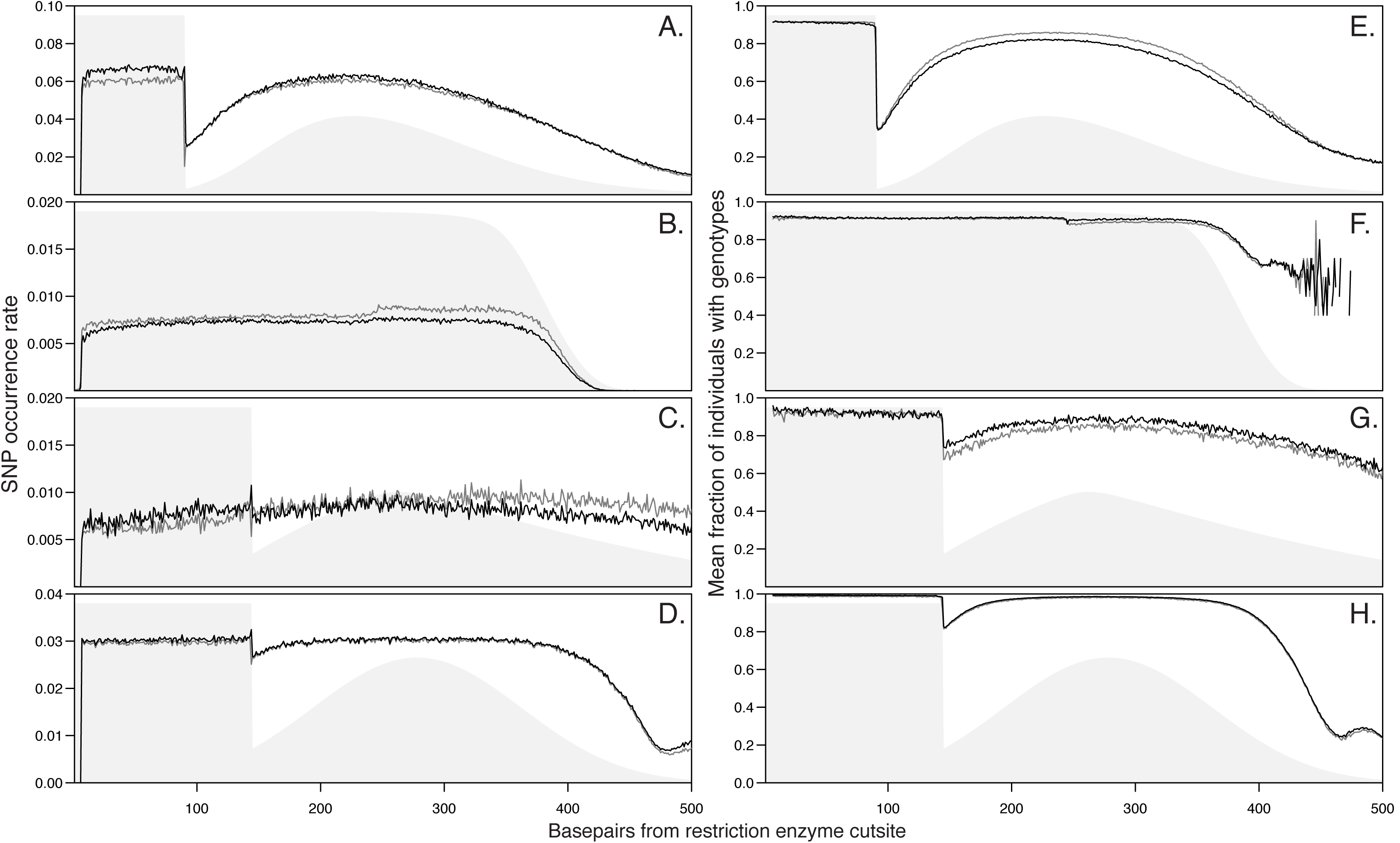
Variation of the average SNP (A-D) and genotype (E-H) call rates along the length RAD loci for each of the benchmark paired-end datasets (A, E: yellow warbler; B, F: threespine stickleback; C, G: bald notothen; D, H: simulations, 20x). The shared areas represent the sequencing coverage underlying the calls (see Fig. 4). Consistent patterns are observed for the reference-based (black lines) and *de novo* (gray lines) approaches across the entire length of RAD loci, demonstrating that Stacks v2 appropriately handles the reverse reads for paired-end data from sdRAD libraries. Coverage is the main driver of the variation of call rates, with genotype calls being more sensitive to reduced coverage than SNP calls.

### Stacks v2 accurately calls and phases genotypes in a reference-based context

The number of SNP and genotype calls may be inflated by erroneous calls, and thus cannot in itself be regarded as an indicator of the quality of these inferences. In order to assess the quality of the Stacks v2 SNP and genotype calls and their spatial distribution within loci, we created simulation datasets with the RADinitio package [Rivera-Colón et al., in prep.], using the threespine stickleback reference genome as template (see Methods). We simulated two datasets, one high coverage (20X) and one low coverage (10X), each comprising 100 individuals split among four populations. Both datasets were based on the same population genetics parameters (see Methods) and had a nucleotide diversity (π) of 0.8%. For simulated datasets, the true loci, read alignments, positions of SNPs and genotypes are known, thus we could calculate the recall and precision of each step, and compare them between a range of methods and approaches (Fig. 5D and 5H, Fig. 6).

**Figure 6.**
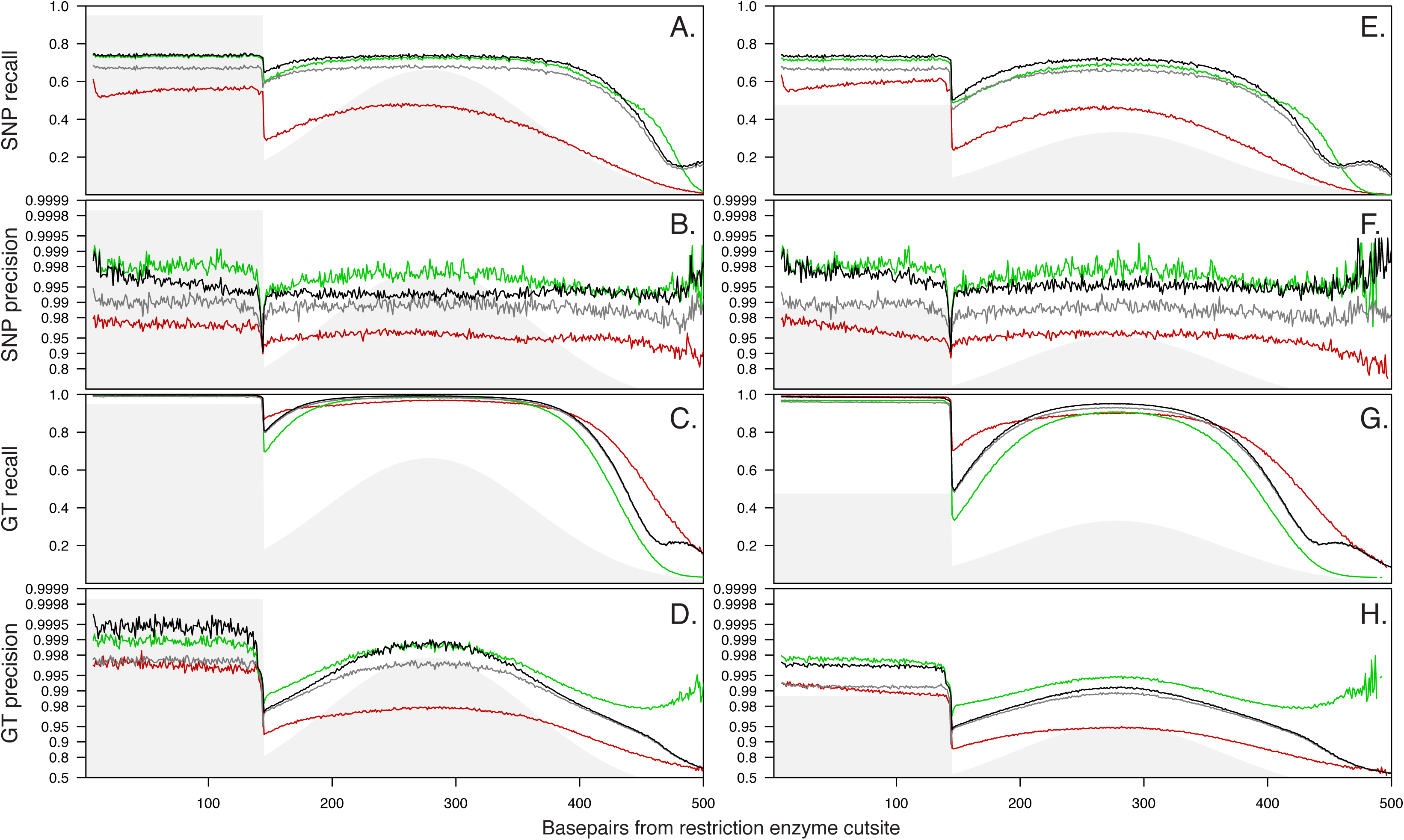
SNP and genotype (GT) precision and recall observed across RAD loci for the 20X (left) and 10X (right) for Stacks v2 using a reference-based approach (black), Stacks v2 using a de novo approach (grey), ANGSD (green), and dDocent (red).

We filtered results for a minor allele count (MAC) of ≥3, because SNP recall and precision correlate with the prevalence of the minor allele of the SNP (expressed as minor allele frequency, MAF, or MAC; Fig. S3). Indeed, SNPs where the minor allele is rare are more likely to be errors or to be missed, but the relationship is non-linear and itself depends on coverage: precision and recall are markedly lower for SNPs with MAC=3, even for well-behaving simulated datasets (Fig. S3). Since our RADinitio software includes a model of the sequencing process (which involves sampling the underlying *true* molecules), even though we know the location of all simulated SNPs, many of them will be ‘sequenced’ at a coverage too low for Stacks v2 to detect them. In this case, a MAC of 3 ensures that an allele is seen in at least two of our diploid samples.

The precision and recall of SNP and genotype calls vary across the length of RAD loci, in tight correlation with coverage, as is shown by our simulations at 20x coverage (Fig. 6A-D) versus 10x (Fig. 6E-H). As noted above, the relationship is more apparent with genotypes than with SNPs, because SNP calls use reads from the entire meta-population and thus depend on the total (rather than per individual) coverage, which is consistently high. SNP precision in the 10x dataset (Fig. 6F, black) was as high as in the 20x one (Fig. 6B, black), and SNP recall was consistent across the locus. Unsurprisingly, genotype recall and precision were lower in the 10x dataset (Fig. 6H versus 6D, black), and deteriorated quickly in regions that are more rarely covered by paired-end reads as a result of the insert size distribution.

We assessed the influence of the read aligner (i.e. BWA alignments) on precision and recall, by comparing results derived from BWA reconstructed alignments with results based on the true, simulated alignments (Fig. S4). SNP and genotype precision rates were nearly identical for both alignment sets, except for a drop at the last few positions of the forward region with reconstructed alignments. This drop is expected and is due to the intrinsic inability of short-read aligners to place indels reliably unless several conserved bases on both sides of the indel are available (no drop occurs on the 5’ end because read alignments are anchored by the conserved restriction site). The near-equal precision at all other positions indicates that read alignments can be reconstructed quite accurately, and contribute little to the error rate or error patterns across the locus.

Next, we compared the inferences of Stacks v2, which implements the [Maruki2017] low-coverage model, with those of ANGSD [Korneliussen2014], using the same BWA read alignments for both methods. ANGSD is a package for SNP and genotype calling that is commonly used for whole genome data, that emphasizes the use of genotype likelihoods and probabilities rather than of explicit genotype calls, but that is nevertheless regarded as more accurate than GATK for the latter [Korneliussen2014, Maruki2017]. For SNP discovery, we found that Stacks had a higher recall (Fig. 6A,E, black versus green), but a fractionally lower precision (Fig. 6B,F, black versus green). We can contrast SNP discovery with calling genotypes. The former occurs in the meta-population, the latter is performed on individuals. When considering the rates of genotype recall and precision it is important to note that only SNPs that were identified correctly can be genotyped, so genotypes can be determined only for sites where a SNP was correctly identified. For calling genotypes the two methods use different statistical approaches: ANGSD uses a posterior probability cutoff, whereas Stacks uses a p-value cutoff. As there is no natural equivalence between the values for these thresholds, we chose the ANGSD cutoff value P≥95% so that the recall was approximately equal to that of Stacks. As a result, the recall was similar for both methods. For genotype precision, the comparison between the two methods depended on coverage. Stacks performed better than ANGSD at higher coverages, and worse at lower coverages (Fig. 6D,H, black versus green).

### Stacks v2 can efficiently cluster RADseq loci, assemble small locus contigs from paired-end reads and map these reads back to loci

To evaluate the ability of Stacks v2 to process sheared, paired-end RADseq datasets when a genome is not available, we performed re-analyses of the datasets introduced above (threespine stickleback, yellow warbler, and bald notothen) using a *de novo* approach (see Methods). We reconstructed respectively 360,292, 89,016, and 34,460 loci present in ≥80% of individuals (NS80 loci, Table 1). Loci present in more than one or two individuals were generally present in most individuals (Fig. S5).

Our *de Bruijn* graph method was able to assemble a consensus contig or a two-contig scaffold for 97.9%, 91.1% and 84.2% of these loci. Although `gstacks` can resolve microsatellite repeats in RAD loci, failure to assemble a consensus contig was essentially due to more complex repeated elements. The contig lengths (mean±standard deviation) were 433±14, 597±59 and 715±47 basepairs (Fig. 7, Table 1).

**Figure 7.**
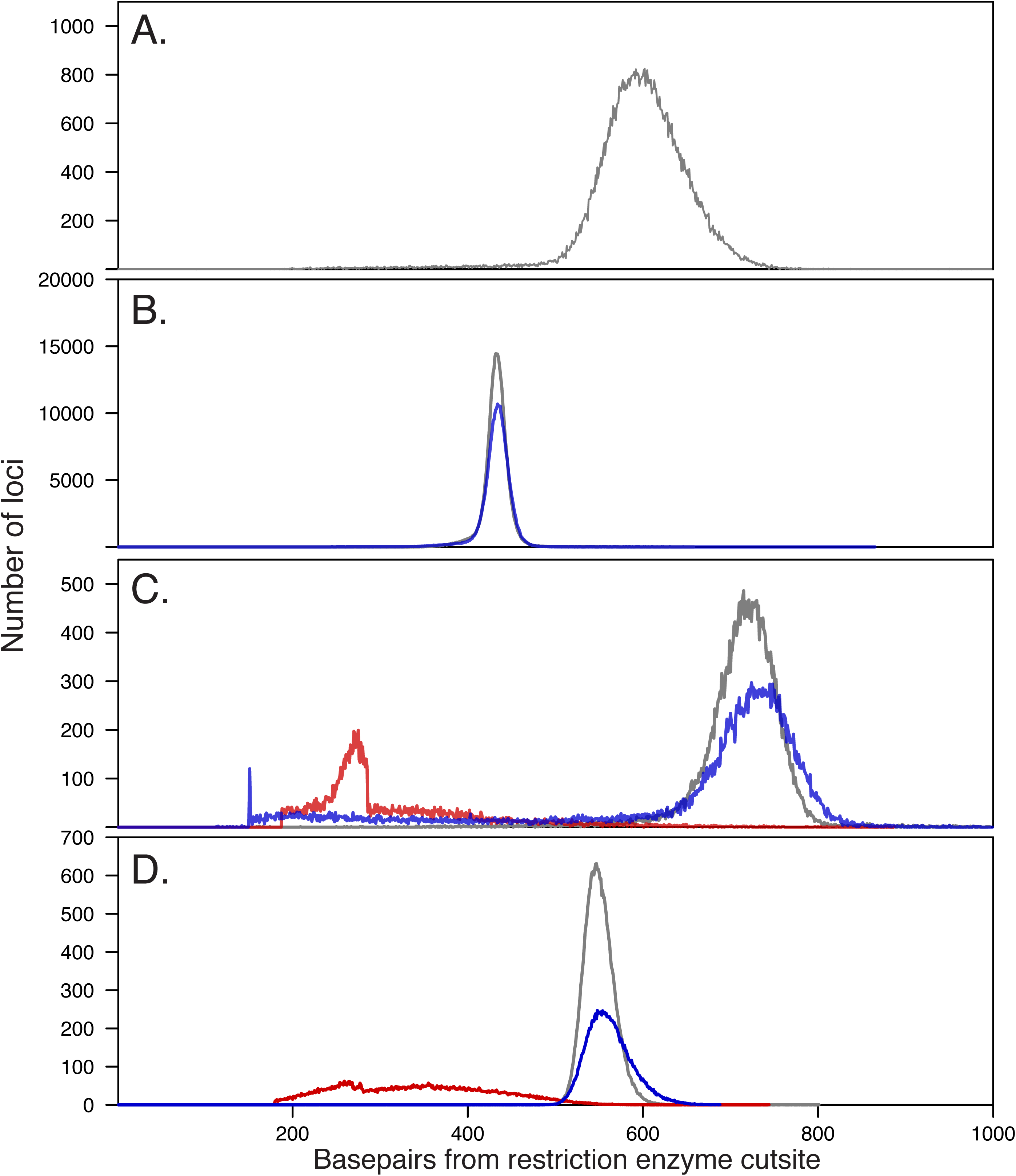
Observed lengths of the RAD loci reconstructed using de novo approaches. by Stacks v2 (gray), RADassembler (blue) and dDocent (red) for the (A) yellow warbler, (B) threespine stickleback, (C) bald notothen, and (D) simulations datasets. Reconstructed locus sizes primarily depend on the shape of the distal tail of the insert size distribution (Fig. 4). The total number of loci represented by each curve corresponds to column XX in Table 1, *i.e.* the plots do not include loci for which the forward and reverse regions could not be combined in a single contig. No data is shown for dDocent in panels A and B because the program did not allow for the analysis of these large datasets (see Methods).

The ability to provide a single, overlapped contig for the entire locus, rather than one contig for the restriction site end and another for the sheared end, depends primarily on the read length and distribution of insert sizes. Indeed, for the threespine stickleback, *P. borchgrevinki*, and yellow warbler datasets, which used 250bp-, 150bp, and 100bp-long reads and insert sizes of 380bp, 309bp, and 392bp (see above), virtually all NS80 loci (>99%) were overlapped.

Once a consensus sequence has been determined for each locus, reads can be mapped back to these loci by our suffix tree-based alignment algorithm. Our method was able to map back 96.1%, 99.3%, and 98.5% of the reads respectively for the threespine stickleback, yellow warbler, and bald notothen datasets (for reads in NS80 loci). We validated our internal read alignment method by comparing the SNPs and genotypes they yielded to those obtained by extracting the consensus locus sequences, aligning reads to them using BWA, and applying the rest of the Stacks v2 pipeline. We found that the results derived from the internal alignments and those derived from the BWA alignments were extremely close in their general statistics, such as the depth of coverage patterns or the number of SNPs and genotypes eventually called (Fig. S4).

Once read alignments have been computed, the *de novo* pipeline continues with the same steps as in the reference-based pipeline, as if the locus contigs derived above formed an ad hoc reference genome.

### The de novo approach performs comparably to the reference-based one

A *de novo* approach involves more steps, each of which may introduce error and decrease the final precision or recall of genotypes. In contrast, the reference-based approach involves fewer steps, so there is less room for error accumulation, although in some cases the reference genome that the analysis relies on may be of unknown quality or distantly related to the system.

We evaluated the global efficiency of the *de novo* approach for analyzing paired-end RADseq data by comparing its results with those of the reference-based approach. We analyzed the three datasets introduced above (threespine stickleback, yellow warbler, and bald notothen) to gain a perspective under real conditions, and we analyzed the high- and low-coverage simulated datasets for a performance comparison in absolute terms, since we know the true underlying states.

For the three real and the two simulated datasets, the *de novo* analysis recovered on average 83% of the same NS80 loci found under the reference-based analysis (respectively 83%, 75%, and 87% for stickleback, warbler, and *P. borchgrevinki*, Table 1). These NS80 loci had comparable coverage, as loci from the *de novo* analysis comprised on average 110% as many aligned reads per individual as the reference-based one (respectively 8.3x, 18.4x, and 22.8x read coverage per locus per individual for stickleback, warbler, and *P. borchgrevinki*, Table 1). For all datasets, the observed insert size distributions were identical for the two approaches (Table 1).

Both approaches found similar densities of SNPs. Indeed, the *de novo* approach identified on average 83% of the same MAC≥3 SNPs per locus (Table 1) that were found using the reference-based approach. Furthermore, the SNP discovery rate varied in the same way within each locus (Fig. 5). The *de novo* approach finds fewer SNPs, particularly in the forward read region, likely because of low-coverage alleles discarded during the assembly of primary and secondary stacks (see [Catchen2011] for details). Furthermore, the *de novo* approach called slightly fewer genotypes than the reference-based one (on average 91% as many at 20x coverage, 89% at 10x; Table 1). The reduction was due to a lower call rate in the distal region, despite coverage being equal with both approaches (Fig. 5). Such a reduction in measured precision at the 3’ end of the locus could be due to some whole RAD loci being discarded by the *de novo* algorithm.

In terms of absolute error rates, which we could compute for the simulated datasets, we found that the *de novo* approach was slightly less precise than the reference-based one, but that the error rates of the two approaches were nevertheless comparable in magnitude, and low in both approaches. In both simulated reference data sets, more than 99% of inferred MAC3 SNPs were true positives. In the *de novo* simulated data, more than 98% of inferred MAC3 SNPs were true positives (Table 1). Importantly, precision was consistent across the length of loci, apart for the expected strong correlation between genotype precision and coverage (Fig. 6). Because error rates were low, SNP and genotype recall rates closely reflected the number of inferred SNPs and genotypes (discussed above). The reduction in precision between reference and *de novo* data is likely due to additional error generated in *de novo* assembly from very low coverage alleles being discarded and a small amount of allele over-merging both occurring in `ustacks`.

### Stacks v2’s methods for paired-end RADseq data perform better than those provided by alternative packages

Next, we compared the results of Stacks on sheared, paired-end RADseq data with those obtained with alternate analysis packages, namely RADassembler [Li2018] and dDocent [Puritz2014] (see Methods). Both packages differ from Stacks in their approach. RADassembler first clusters single-end loci using Stacks v1, then builds paired-end contigs with the CAP3 overlap-layout method [Huang1999]. RADassembler focuses only on locus construction and does not identify SNPs or call genotypes. dDocent relies on CD-HIT [Li2006], Rainbow [Chong2012], and PEAR [Zhang2014] for locus and contig assembly and it applies the FreeBayes [Garrison2012] software for SNP identification and genotyping.

In the simulated data sets, the number of NS80 loci RADassembler constructed was approximately 70% of the number of loci constructed in the same *de novo* Stacks analysis. For those loci constructed, the size and number that could be overlapped with the forward locus was very similar to Stacks v2. However, RADassembler reconstructed markedly shorter contigs for a small subset of loci. The dDocent software constructed ∼105% of the loci that *de novo* Stacks v2 did, however, those loci were 70% as long while only 40-45% could be overlapped indicating assembly errors (Figure 7 and Table 1). In the empirical *P. borchgrevinki* data set, RADassembler performed as well as Stacks v2, dDocent, however, was not able to complete processing the data set.

### Stacks v2 is able to provide consistent and rich haplotypes

Producing useable haplotype markers involves two main processes. First, a consistent phasing of SNPs at each locus must be found in each individual. When considering a locus at the population level, however, the haplotype graphs between individuals may be different due to the presence or absence of particular SNPs in particular individuals (see methods). Stacks v2 is able to produce consistent, population-wide haplotypes that incorporate 95% of independently genotyped SNPs in the 20x simulated dataset, and 86% given 10x sequencing. For the empirical datasets, the value ranged from 71-97%. The haplotypes built were nearly 10bp in mean length in the yellow warbler, versus approximately 3bp in stickleback and the bald notothen, suggesting the warbler genome is highly polymorphic.

## Discussion

### Long RADseq contigs

The appeal of using paired-end sequencing to derive short contigs from *de novo* sdRAD data has been promoted repeatedly [Etter2011; Amores2011; Hohenlohe2013; Andrews2016; Nelson2018; Li2018]. However, the approach has remained elusive due to the technical difficulty, unreliability and inefficiency of generating these contigs in the absence of any software capable of performing this task natively. Thus, [Etter2011] developed an approach combining Stacks and Velvet [Zerbino2008], while [Hohenlohe2013] experimented with both Stacks and Velvet or Stacks and CAP3 [Huang1999], opting for the latter, and [Nelson2018] used Stacks and Fastq-Join [Aronesty2011] together with a longer-read, high-overlap sequencing strategy. Finally, the recently published RADassembler method wraps around Stacks v1 and CAP3 [Li2018] in a similar manner to [Hohenlohe2013].

The series of methodological developments that we introduce in Stacks v2 makes the analysis of paired-end sdRAD data efficient, reliable, and accessible to the majority of RADseq users. It yields more robust results and is considerably faster and easier to apply in comparison with previous approaches. Importantly, the length of the assembled paired-end contigs and the merging rate of the ‘forward’ and ‘reverse’ regions of the locus primarily depend on the distributions of insert sizes in the sequenced DNA library (see below). However, the average contig length can be expected to be in the 400-800bp range, and we find that virtually all loci have a single contiguous contig provided that at least a small fraction of reads overlap.

In the *de novo* context, the availability of RAD locus contigs offers several clear advantages over the shorter loci obtained from single-end data, which are typically only as long as the individual reads. These loci can be more easily mapped to existing genomic resources for the purposes of providing functional annotation, conducting linkage-based genomic scans [Amores2011; Feulner2019], or designing capture baits [Ali2016]. They allow one to make use of the paired-end reads derived from the sdRAD [Baird2008] and BestRAD [Ali2016] protocols, even when a *de novo* strategy is employed. As these protocols involve a random shearing step, this implies that PCR duplicates can be filtered natively based on the start position of the reverse read in the same way as can be done when paired alignments have been made to a reference genome [DePristo2011].

Finally, in addition to the qualitative advantages presented above, using paired-end reads of course increases the amount of genotype data that is produced. Although paired-end data do not change the number of RAD loci, they can more than double the average number of SNPs per locus for suitable library preparation and sequencing parameters (Table 1 and see below). On average, this results in more polymorphic loci, that are each more informative. In a *de novo* context, these results will benefit any study focused on species with low diversity, and in particular phylogenetics studies, where sparse genotype matrices are being constructed across many species [Near2018]. When RAD loci are ordered onto chromosomes (*i.e.* for analyses that use a reference-based approach, or if *de novo*-assembled loci have been mapped to an external reference genome), this amounts to a higher information density along the genome, which should help resolve linkage patterns at a finer scale and identify evolutionary events that may have been missed with a sparser sampling of the genome [McKinney2017].

### Haplotypes

Expecting multiple SNPs at each locus opens the possibility of treating RAD loci as a set of haplotypes rather than as individual SNPs. Depending on the nucleotide diversity of the species, it not rare to see up to ten SNPs per locus in real datasets (Table 1). Since a polymorphic site can have no more than two allelic states, it can only label two alleles in the population. Within a small genomic region, choosing one SNP over another may result in a different evolutionary signal. By phasing these SNPs, Stacks v2 can instead provide multiallelic haplotypes which can encode a much larger amount of information regarding the provenance of the genomic region. These rich, physically phased haplotypes produced by Stacks v2 can be used to study changes in coalescence in different regions of the genome [Nelson2018].

The haplotyping algorithm implemented in Stacks v2 relies strictly on the co-observation of alleles with a read (or read pair). This is in contrast with statistical phasing methods in which an individual’s haplotypes are estimated in relation to a panel of haplotypes observed at the population level [Browning2011]. Furthermore, phasing algorithms that rely on the co-observation of alleles in reads are designed to find the haplotypes that are most consistent with the data, and subsequently accept the optimal haplotypes as the real ones [Patterson2015]. Our approach instead expects the data to appear consistent at a specified tolerance level (see Methods). While this tolerance threshold makes our approach more sensitive to insufficient coverage, it allows one to identify and remove cases where the allelic observations for an individual at a particular locus are at odds with diploidy. Such cases point to miscalled genotypes that can then be pruned out, or occasionally to contamination in the sequencing library (that makes an ‘individual’ effectively non-diploid) as evidenced by the dramatically reduced phasing rates that may be observed for specific individuals (Fig. S6). Over-merged loci, that collapse several paralogous genomic loci into a single polyploid one, also exhibit high phasing failure rates and can be filtered on this basis.

### Stacks v2 has lots of improvements

Stacks v2 supports nearly all major protocols for reduced representation, marker-based experiments, including single- and double-digest RAD [Peterson2012] and DaRT [Kilian2012], using single- or paired-end sequencing, as well as 2bRAD [Wang2012] and GBS using single-end sequencing. We note that it is not suitable for paired-end data derived from GBS [Elshire2011], as forward and reverse reads contain essentially the same information. Applying our approach to paired-end GBS data would lead to assembling two half-coverage copies of each locus (one for each possible orientation).

One of the major advantages to the design of Stacks v2 is the vertically integrated nature of the software system. This design ensures that each stage of the analysis has access to all of the information collected so far, from the results of single-end clustering, to the resulting *de Bruijn* graph fragments assembled, or the matrix of genotypes. Most importantly, Stacks can take advantage of certain types of information that RADseq data provide, for example, the *de Bruijn* graph assembler knows that all reads in a RADseq analysis are sequenced in the same direction (and hence the *de Bruijn* graph only has to account for one strand). This is in contrast to the majority of alternative software pipelines available (two of which we compared to Stacks v2), where they incorporate existing software in a black box way (that is, how the software operates is opaque to the executing pipeline), and where such software was not designed specifically for RAD data. In many cases, we could not run competing software (dDocent and RADassembler) because the softwares were unable to accommodate the large (but quite common in size) data sets we tested above.

### Efficiently applying paired-end RADseq

The particular nature of RADseq libraries has important implications with regard to the distribution of coverage across the locus. To achieve an optimal insert size distribution, we recommend that the width of the size selection window during library construction range between two and three (or less than three) read lengths (plus the length of the adapters). So, for 150bp reads, an optimal insert length distribution could be obtained by size selecting for 300-400bp inserts (that is, for 430-530bp molecules if using 130bp adapters). In addition, overlapping of the forward and reverse regions into a single contig is only possible when there is overlapping within at least some read pairs. In practice, if at least 5% of reads overlap (that is, have inserts shorter than two read lengths plus adapters), high overlapping rates will be observed.

### RADseq PCR duplicate rates must be monitored

Nearly all RAD protocols use PCR amplification as a method to enrich DNA adjacent to restriction enzyme cut sites. Some protocols, in fact, use PCR as a method to enrich shorter molecules adjacent to cut sites over longer molecules in lieu of an explicit size-selection of molecules using beads or a gel. However, PCR amplification can create an illusion as to how much genetic information is available in a RAD library, particularly if the starting amount of template DNA was very small, and can result in bias when some alleles or loci are not sampled. In addition, the amplification process introduces sequence errors (on the order of 10^−7^ per nucleotide per cycle for the commonly used Phusion enzyme [NEBiolabs; https://www.neb.com/faqs/2012/09/06/what-is-the-error-rate-of-phusion-reg-high-fidelity-dna-polymerase]) and the accumulated error from many rounds of PCR can introduce sites that will falsely appear polymorphic in small numbers of individuals.

In our analysis, substantial PCR duplicate rates were observed in all datasets, ranging from 33% (yellow warbler) to 76% (bald notothen, Table 1). We do not doubt these figures, as insert sizes were reconstructed robustly (see Results), and as we could confirm them with alternative software in the case of reference-based analyses. The rates differed somewhat between individuals within datasets, but most of the differences occurred between datasets or possibly between libraries (Fig. S7). The BestRAD [Ali2016] protocol (for which we only had two datasets from the same group) seemed to yield fewer PCR duplicates (33%) than the original [Baird2008] protocol (41-76%). This makes sense, as the BestRAD protocol is the only RAD protocol that uses biotinylated adapters so that the restriction site-adjacent DNA can be extracted from the remaining genomic DNA using beads early on in the library preparation.

For ddRAD protocols, it is not usually possible to identify PCR duplicates, but methods based on degenerate adapters have been developed. [Schweyen2014] have reported PCR duplicate rates of 20-45%, and [Tin2015] have reported rates of 30-80%. While these figures represent varying systems, preservation states and protocols, they suggest that PCR duplicates are as prevalent in ddRAD as in shearing-based RAD protocols.

PCR duplicates are believed to interfere with genotype calls—in particular, to reduce the power to call heterozygotes by displacing the allelic ratio—and thus to bias downstream analyses. We note, however, that little effort has been made to accurately measure the impact of PCR duplicates. Such work would be useful both to confirm the theoretical argument that PCR duplicates alter genotype calls, and to estimate the extent of the bias that approaches that do not allow PCR duplicate removal putatively suffer from. Nevertheless, we encourage RADseq users to use protocols that readily allow the removal of PCR duplicates.

### Choosing the right protocol to maximize analytical results

With hindsight, we have chosen to focus our software development on the most effective molecular protocols. In general, sdRAD ([Baird2008; Ali2016]) experiences less allelic dropout or bias than ddRAD ([Peterson2012]), and specifically, BestRAD ([Ali2016]) improves analytic possibilities among sdRAD protocols through its method of separating RAD DNA from genomic DNA early in the molecular protocol. Our data implies that the rate of PCR duplicates coupled with sequencing coverage is the best measure of library quality and the best predictor of analytical results. Experimenters should measure PCR duplicates (which requires the use of sdRAD or the 3RAD [Graham2015] protocol) and ensure robust PCR amplification did not mask low amounts of template DNA and they should ensure, that after filtering PCR duplicates from their data, that they still have reasonable sequencing coverage for analytical purposes.

## Conclusion

The family of RAD protocols developed over the past decade, coupled with commodity-priced, massively parallel, short-read sequencing, has found a valuable niche for conducting large population genomic and phylogenomic studies. We can optimize the accuracy and volume of data available to researchers employing reduced-representation strategies by providing a set of accurate, fast and accessible software. Stacks v2 provides the analytical tools to enhance RADseq when it is coupled with paired-end sequencing, assembling tens of thousands of loci that are 400-800bp in length and can contain up to a dozen physically-phased SNPs on each locus. The software can scale to datasets with thousands of individual samples and we have shown here the effectiveness of the algorithms to assemble loci and genotype those individuals. By focusing software changes to provide robust genetic information in the current sequencing landscape we can fortify the utility of RAD sequencing for another decade.

## Acknowledgements

The authors would like to thank Thom Nelson, Bill Cresko, and Susan Bassham for useful discussions and Rachel Bay for help accessing the yellow warbler data. We would also like to thank the users of Stacks for all of their input, bug reports, and early testing of the software. AGR and NCR were supported by NSF grant XXXX.

## Supplementary Figures

**Figure S1**

(A) Comparison between the insert length distribution in the P. borchgrevinki RADseq library reconstructed from read alignments by gstacks (solid line) and the digitalized version (dotted line) of the (B) raw output of the fragment analysis for this DNA library.

**Figure S2**

SNP and genotype (GT) precision and recall plotted as a function of the average per-sample coverage at the locus position, for the reference-based (black) and *de novo* (red) Stacks analyses of the the 20X (diamonds) and 10X (crosses) simulated datasets. The data corresponds to that of Fig. 6 of the main text.

**Figure S3**

Properties of the MAC<3 SNPs. (A) Precision across the locus for MAC=1 SNPs for the reference-based Stacks analysis of the 10X simulated dataset; compare with Fig. 6F of the main text, that uses MAC≥3 SNPs. (B) Average precision at positions 25-124 as a function of the (inferred) minor allele count. (C) Recall at positions 25-124 as a function of the minimum inferred MAC threshold (*e.g.* recall is ∼0.7 when considering MAC≥3 SNPs).

**Figure S4**

Comparison between the SNPs obtained with the native Stacks v2 *de novo* read aligner and those obtained by substituting BWA into the *de novo* Stacks v2 pipeline for the warbler (top), notothen (center) and 20X simulated (bottom) datasets.

**Figure S5**

Histograms of the number of individuals per locus, for each dataset (columns) and according to the Stacks reference-based (top) or de novo (center) approaches or to dDocent (bottom). No plots are shown for dDocent for the warbler and stickleback datasets as the program did not complete.

**Figure S6**

Phasing failure rate for each of the 241 individuals of the warbler datasets. Elevated phasing failure rates in some individuals (top) suggest quality issues for these samples, such as contamination.

**Figure S7**

Boxplot of the per-individual PCR duplicates rates observed within each library (the 241-individual warbler dataset is composed of three libraries).

**Figure S8**

Insert length periodicity is 10.5bp for warbler, the largest dataset. This length seems to match the 10.5-basepair period of the DNA helix, which suggests that this observation may be caused by a physical process, possibly by the shearing step.

